# Detecting Drug-Target Binding in Cells using Fluorescence Activated Cell Sorting Coupled with Mass Spectrometry Analysis

**DOI:** 10.1101/121988

**Authors:** Kris Wilson, Scott P Webster, John P Iredale, Xiaozhong Zheng, Natalie Z Homer, Nhan Pham, Manfred Auer, Damian J Mole

## Abstract

The assessment of drug-target engagement for determining the efficacy of a compound inside cells remains challenging, particularly for difficult target proteins. Existing techniques are more suited to soluble protein targets. Difficult target proteins include those with challenging *in vitro* solubility, stability or purification properties that preclude target isolation. Here, we report a novel technique that measures intracellular compound-target complex formation, as well as cellular permeability, specificity and cytotoxicity - the Toxicity-Affinity-Permeability-Selectivity (TAPS) technique. The TAPS assay is exemplified here using human kynurenine 3-monooxygenase (KMO), a challenging intracellular membrane protein target of significant current interest. TAPS confirmed target binding of known KMO inhibitors inside cells. We conclude that the TAPS assay can be used to facilitate intracellular hit validation on most, if not all intracellular drug targets.

## Introduction

The efficacy of a drug relies upon interaction with a relevant therapeutic target protein at the physiological site of activity. Direct detection of this interaction *in vitro* remains a challenge, particularly if isolated protein cannot be obtained or if a suitable assay method cannot be designed. The measurement of binding of a hit, lead or drug to its intended target is particularly challenging in cells. Therefore, detection of target engagement is one of the major challenges in hit validation from phenotypic assays ^[1-3]^. Several methods have been reported for detection of drug-target engagement. A selection of creative techniques which utilise functionalised drugs or probes with fluorescence polarized imaging have recently been described ^[4-7]^. Others use chemoproteomic methods to profile the spectrum of proteins interacting with chemical probes ^[8]^. Such techniques require specialised/adapted instruments and/or capacity for production of fluorescently labelled probes. The current standard for demonstrating intracellular target engagement is the cellular thermal shift assay (CETSA), which detects ligand-induced thermal stabilisation of target proteins ^[9-11]^. However, the authors state that the method is “not likely to work for highly inhomogeneous proteins”. In general, challenging drug targets which are not readily expressed in a soluble form, which are unstable in solution or which behave differently in the absence of key co-factors or protein binding partners pose difficulties. We developed the TAPS method to provide a solution for this gap in target engagement measurement and to extend the toolbox of approaches for soluble protein targets. TAPS combines fluorescence activated cell sorting (FACS) with liquid chromatography mass spectrometry (LC-MS/MS) analysis to detect binding of drugs to their target inside cells in a concentration-dependent manner producing results which resemble an apparent binding curve of a small molecule to a target in any cellular context. Additionally, TAPS selects, in a single assay method, compounds with sufficient cellular permeability, and excludes, via necessary controls, compounds which lack specificity or demonstrate high cytotoxicity. This method is generally applicable for any intracellular target which can be expressed as a fluorescent fusion protein and its applicability to challenging membrane proteins is demonstrated here.

The TAPS technique involves transient expression of the gene of interest as a fluorescent protein conjugate in an appropriate cell type to generate a range of target concentrations. Thus, fluorescently tagged target proteins are assayed in a cellular environment in the presence of endogenous co-factors and protein-protein interactors in a physiologically relevant tissue. The protein is produced within 24 hours and the transfected cells are incubated with single or pooled compounds. Cells are subjected to FACS analysis and sorted according to fluorescence intensity exhibited by the fluorescent fusion protein. Lysis of the sorted populations is followed by liquid chromatography mass spectrometry (LC-MS/MS) analysis, allowing comparison of compounds bound in target-free non-transfected cells and those bound in the fluorescently intense cells expressing varying levels of the target protein. Negative cells resulting from transient expression in these experiments act as an important integral control allowing the specificity of drug binding to be determined.

The TAPS assay was developed using the mitochondrial membrane associated enzyme kynurenine 3-monooxygenase (KMO). This enzyme is emerging as an increasingly important target for drug development since it has recently been implicated as a therapeutic target for Huntington’s disease ^[12]^ and multiple organ failure caused by severe acute pancreatitis ^[13]^. However, human KMO is difficult to produce and isolate in a recombinant form since it is membrane-associated and requires the presence of a lipid and proteinaceous environment to maintain activity. The hydrophobic membrane-targeting domain at the C-terminus of this NADPH-dependant flavoprotein hydroxylase is believed to be responsible for its low aqueous solubility, poor stability, and tendency to aggregate with other membrane proteins when expressed recombinantly ^[14-16]^. Therefore, the importance of assaying drug-KMO engagement directly in mammalian cells in the presence of co-factors and intracellular binding proteins required for enzyme functionality was paramount. Development and successful application of the TAPS method for assaying compounds against KMO inside cells provides a strong indication that this technique is applicable for targeting other challenging proteins.

## Materials and Methods

### Cloning

The E2-Crimson-human KMO gene (Cys452 variant) was synthesised by GenScript in vector pUC57. The DNA sequence for E2-crimson was sourced from Clontech. The gene for the fluorescent-KMO fusion was ligated into vector pCDNA3.1 (Invitrogen) using restriction sites NheI (N-terminus) and NotI (C-terminus).

### Transient Expression of Fluorescent Target Protein

HEK293 cells were passaged in poly*-D*-Lysine treated plates and incubated overnight in OPTI-MEM medium (Lonza) at 37°C, 5% CO_2_. The cells were transiently transfected the following day with pcDNA3.1-E2-Crimson-huKMO DNA using Lipofectamine 2000 (Invitrogen) in OPTI-MEM medium by standard transfection protocol. Transfection medium was removed from the cells 6 hours post-transfection and replaced with DMEM containing 10% FBS, 1% L-glutamine and 1% penicillin-streptomycin (Life Technologies). The transfected cells were maintained at 37 °C, 5% CO_2_ for 24-48 hours post-transfection before the TAPS assay was performed.

### TAPS Assay

#### Step 1: Compound Incubation

Compounds were diluted to a concentration of 20 μM (DMSO <1%) in tissue culture medium (DMEM with 10% FBS, 1% L-Glutamine, 1% penicillin-streptomycin), then incubated with the transfected cells at 37 °C, 5% CO2. Following incubation, the cells were detached from the plate by gentle pipetting and centrifuged for 5 minutes at 1000 rpm. Culture medium was removed by pipetting and the cell pellet re-suspended in FACS buffer (PBS + 2% FBS) to wash off unbound compound. The cells were centrifuged, as above, and the wash buffer removed. The cells were then re-suspended in FACS buffer and transferred to 5 mL FACS tubes. The tubes were wrapped in foil and placed on ice until sorting.

#### Step 2: FACS Sorting

Cells were sorted using a BD FACS Aria II system fitted with a 100 micron nozzle. Data was acquired and processed using BD FACSDiva Software version 6.1.3. Cell fluorescence was detected using the APC channel, using the 640 nm laser at 40 mWatt for E2-Crimson excitation. The filter used was 670/14 nm detecting fluorescence emission of E2-Crimson in the 663 – 677 nm range. Cells were sorted and collected into four cell populations defined by fluorescence intensity, forward and side scatter gating was used to exclude dead cells and only live cells were collected. Cells were collected in 5 mL FACS tubes containing 1 mL of DMEM with 10% FBS, 1% L-Glutamine and 1% penicillin-streptomycin. Each population of cells was centrifuged at 1000 rpm for 5 minutes. Medium was removed and the cell pellet re-suspended in 20 mM HEPES, pH 7.0. The suspension was briefly sonicated to lyse the cells before centrifugation, as before, to pellet cell debris. The lysate (supernatant) was transferred to LC-MS vials and stored at -20 °C prior to MS analysis.

#### Step 3: Mass Spectrometry Detection of Compounds in Cell Lysates

##### Instrumentation

This method was previously described by us ^[17]^. The chromatographic system used was a TurboFlow^™^ TLX Aria-1 system (Thermo Fisher Scientific, Hemel Hempstead, UK), consisting of two Allegros pumps defined as the loading and eluting pumps, two valve-switching modules and a CTC liquid autosampler. The detection was carried out using a TSQ Quantum Discovery triple quadrupole mass spectrometer (Thermo Fisher Scientific, Hemel Hempstead, UK).

##### Chromatography and Mass Spectrometry parameter optimisation

Samples were subject to online-extraction using a TurboFlow^™^ TLX Aria-1 system operated in focus mode. 10 μL injection of the cell lysate was loaded directly onto a C18PXL (50 x 0.5 mm, ThermoFisher Scientific, Hemel Hempstead, UK) TurboFlow^™^ column at a high flow rate, causing the proteinaceous material to flow to waste. A series of valve switches led to the elution of the extracted sample from the TurboFlow^™^ column directly onto the analytical column. Solvent A was water with 0.1% formic acid, Solvent B was methanol with 0.1% formic acid and Solvent C was 45:45:10 acetonitrile:isopropanol:acetone.

Following TurboFlow^™^ extraction, the analytes were subsequently separated on a reverse phase T3 Atlantis (2.1 × 150 mm, 3 μm, Waters, Manchester, UK) analytical column, protected by a Kinetex KrudKatcher^®^ (Phenomenex, Macclesfield, UK). The analytical column was maintained at 5 °C using a column chiller.

The online TurboFlow system Aria-1 was directly connected to a Quantum Discovery triple quadrupole mass spectrometer, operated in electrospray ion mode with polarity switching for positive and negative ion monitoring. The source temperature was 300 °C, the spray voltage 3 kV and the skimmer offset was 12 V. Argon, the collision gas in Q2, had a pressure of 1.5 mTorr. Automated tune settings were used to achieve the maximum ion signal for each analyte for initial validation experiments, optimizing on tube lens voltage, parent to product transitions and collision energy for each transition. A quantifier and qualifier ion was determined for each analyte. By monitoring for quantifier and qualifier ions this adds additional specificity to the assay. Acceptable quantifier: qualifier peak area ratios in biological samples were considered to be those that fell within 20% of the average ratio seen in standards. By applying tune settings, a peak area could be generated for each compound. For pooled compound screening, tune settings were not used. Samples were subject to a full MS scan with the molecular weight detection range set at 250 to 465 (Daltons). Peaks detected in this initial scan were then identified by molecular weight and checked versus their mass-charge ratio.

A scan width of m/z 0.5, scan time of 0.1 s and unit resolution on Q1 and Q3 were applied. Data was collected as centroid data to minimize the file sizes. Data were acquired and processed using Aria 1.3, Xcalibur 1.4 and LC Quan 2.0 SP1 software packages.

##### Compounds

Three known KMO inhibitors from the patent and scientific literature ^[18, 19]^ and a nonbinding compound from the University of Edinburgh Drug Discovery Core compound library were used for assay development and validation (Figure 1). 100 compounds selected from the University of Edinburgh Drug Discovery Core compound library were pooled with compounds 1, 2 and 3 (Figure 1) and assayed simultaneously as a mixture to demonstrate multi-compound screening.

#### KMO Activity Assay

All compounds assayed in the compound “pool” were evaluated for activity against human KMO in a KMO activity assay. The source of KMO protein for this assay was lysate generated using the following stable cell line:

HEK293 (Flp-In-293 which express lacZ-Zeocin, Life Technologies) cells were stably co-transfected with 9 μg of pcDNA5/FRT/V5/HisTOPO DNA and 18 μg of pog44 plasmid. Pog44 expresses Flp recombinase protein, co-transfection of pog44 with the gene of interest allows targeted integration into the mammalian cell genome within a transcriptionally active region. Transfection was carried out using Lipofectamine 2000 (Life Technologies) in OPTI-MEM medium (Lonza). Cells were selected for two weeks using Hygromycin B (Sigma Aldrich) at 100 μg/mL to select for the hygromycin resistance gene contained within the pcDNA5 vector before positive colonies were isolated and cultured.

HEK-huKMO cell lysate was analysed for KMO enzymatic activity by measuring the amount of KYN converted to 3HK detected using liquid chromatography-mass spectrometry (LC-MS/MS) following incubation with the compounds screened in the TAPS assay using a method we described previously ^[17]^. Compounds were tested in a one compound per well format in this assay. Briefly, lysate containing 200 μg total protein was incubated with 4 mM MgCl2, 1 mM NADPH and 200 μM L-Kynurenine in 20 mM HEPES, pH 7.0 for two hours at 37°C with gentle shaking at 250 rpm in a total assay volume of 100 μL. Samples were added to 500 μL acetonitrile (containing 25 μg/mL of internal standard, d5-TRP) to terminate activity, followed by centrifugation at 4000 rpm for 20 minutes to pellet the precipitate. The supernatant fraction was dried under nitrogen, and the residue re-suspended in 30:70 methanol:water with 0.1% formic acid ready for LC-MS/MS analysis.

LC-MS analysis was carried out using the TSQ Quantum Discovery triple quadrupole mass spectrometer (Thermo Fisher Scientific, Hemel Hempstead, UK) using a pentafluorophenyl (PFP) fused pore column (Waters), at 40°C. The injection volume was 10 μL and the flow rate was 500 μL/minute. The method had a run time of 4 minutes and d5 Tryptophan was used as an internal standard. Qualifier and quantifier peaks were identified for 3HK and for d5 Tryptophan. Data was acquired and processed using Xcalibur 1.4 and LC Quan 2.0 SP1 software packages.

#### Cell Staining

HEK293 cells were transiently transfected and sorted by FACS as described above. A sample of cells from each sorted population were plated in transparent 96 well plates (Whatman) and incubated overnight at 37°C, 5% CO_2_. Medium was aspirated from the cells the following day and the cells gently rinsed with PBS^+^ (PBS containing 1 mM Ca^2+^ and 0.5 mM MgCl2). Wheat germ agglutinin-AlexaFluor488 conjugate was used at a concentration of 5 μg/mL to stain the cell membranes by incubation for 10 minutes at 37 °C. The stain was removed and the cells rinsed twice with PBS^+^. The cells were then fixed by ten minute incubation in 3.7% formaldehyde diluted in PBS^+^. After removal of the fixative, cells were thoroughly rinsed three times by five minute incubation in PBS^+^. To permeabilise the cells, 0.1% Triton-X 100 in PBS was added for 5 minutes at room temperature. Cells were washed a further three times before incubation with 300 nM DAPI in PBS for 5 minutes at room temperature in the dark to allow nuclear staining. Cells were washed a final three times in PBS. The cells were imaged using the Opera™ High Content screening system. E2-Crimson-huKMO fluorescence was detected using the 640 nm laser (2000 μW, 280 ms), nuclear (DAPI) staining was detected using the UV light source (365 nm excitation, emission filter 450/50, 40 ms) and the 488 nm laser (1250 μW, 40 ms) was used to detect the cell membrane stain (wheat germ agglutinin-Alexa488 conjugate).

## Results

### Selection and Validation of KMO Inhibitors

Three known inhibitors of KMO, with nanomolar to micromolar affinity^[18,19]^, and one structurally related, negative control compound (Figure 1) were selected for development and validation of the TAPS assay. The activity of these inhibitors was verified by us previously against human KMO ^[17]^ in the enzyme activity assay described in the methods section and the *IC_50_* values generated were in a similar range to literature values reported against rat KMO enzyme for compounds 1-3 (Figure 1). Compound 4 was confirmed to be inactive in the activity assay (Figure 1).

**Figure 1.**
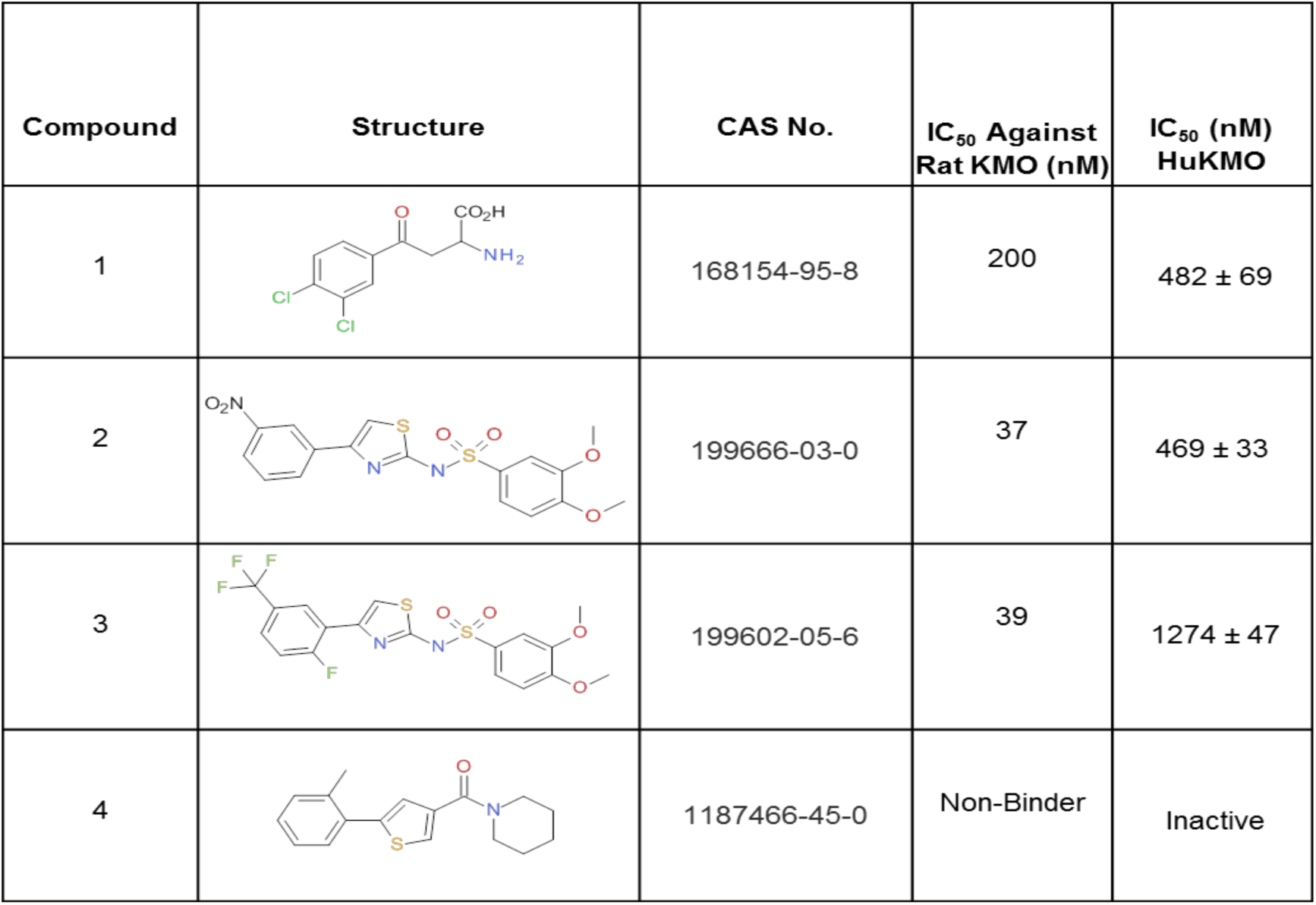
The structures and activities of compounds used to validate the cellular TAPS method. CAS numbers and literature reported activities for rat KMO enzyme are shown for Cpds 1 – 4 alongside the *IC*_*50*_ values generated by us previously in a KMO activity assay against human enzyme ^[17]^.

### Cellular E2-Crimson Fluorescence Correlates with KMO Activity

KMO was detectable in the cellular matrix as a far red fluorescent conjugate by fusion with E2-Crimson^[20]^. A broad range of cellular fluorescence intensities resulting from transient expression of fluorescent KMO were detected and sorted by FACS to generate the cell populations for MS analysis. Confocal imaging of cells from the sorted populations was used as a parallel method to confirm KMO expression (Figure 2a-b). Sorted cell populations, when lysed and assayed for KMO enzymatic activity, showed good correlation between increased fluorescence intensity and KMO enzyme activity (Figure 2c). Therefore, cellular E2-Crimson fluorescence reliably indicated KMO expression and functionality.

**Figure 2.**
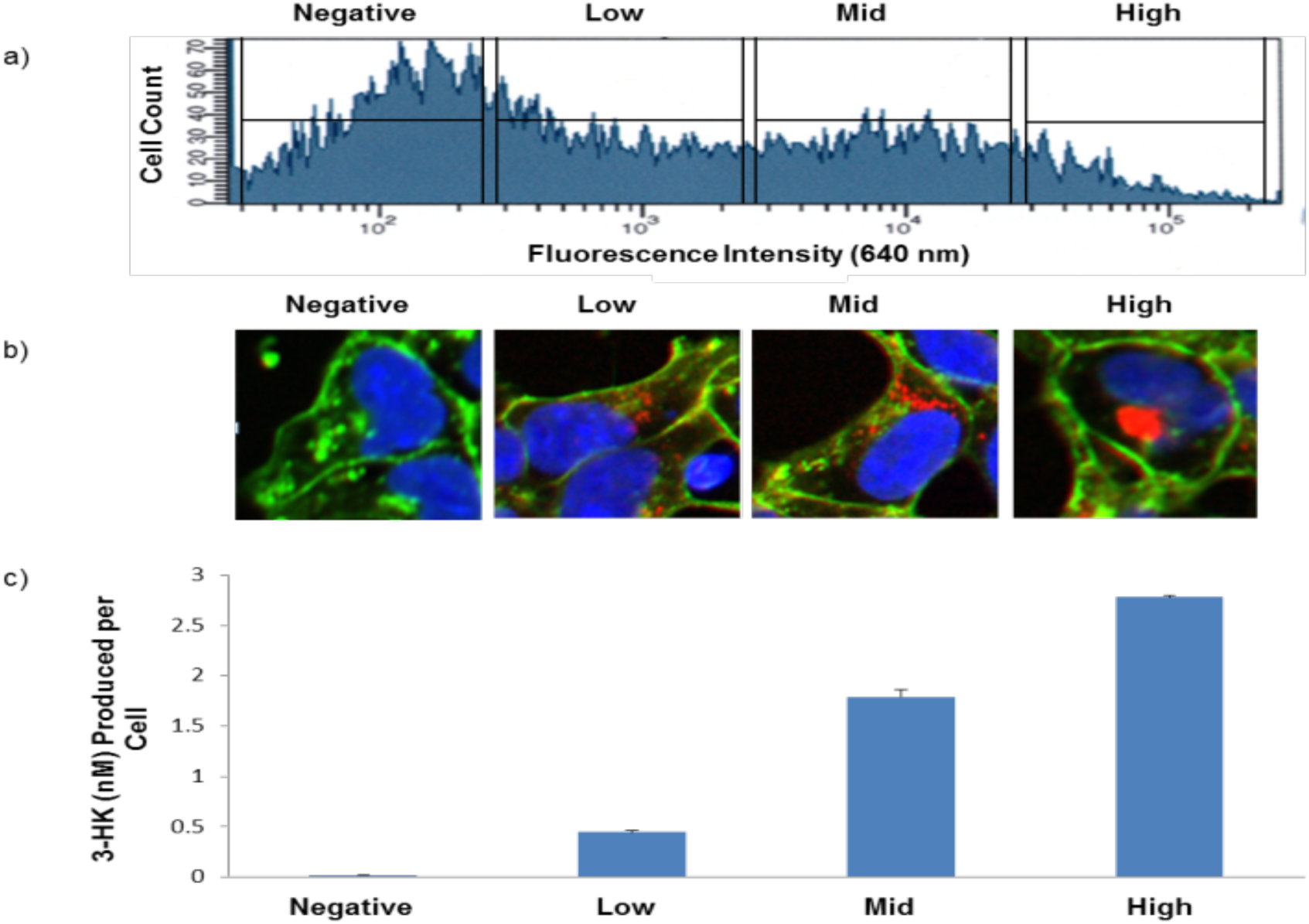
Expression of functional hKMO and compound screening in the TAPS assay. a) Transient transfection of HEK293 cells with E2-Crimson-huKMO generates populations of variably transfected cells, the histogram shows the distribution of fluorescence intensity in the transfected cells and the population gates applied for cell sorting, b) Cellular staining images obtained using the Opera™ High Content Screening system show variable expression of E2-Crimson in each sorted population. E2-Crimson-huKMO fluorescence was detected using 640 nm laser excitation (2000 μW, 280 ms), DAPI staining was detected using the UV light source (365 nm excitation, emission filter 450/50, 40 ms) and the 488 nm laser (1250 μW, 40 ms) was used for exciation the cell membrane stain (wheat germ agglutinin-Alexa488 conjugate). c) 3-HK produced per cell in each sorted cell population lysate after incubation with 200 μM kynurenine substrate for 2 hours correlates with fluorescence intensity in the cell.

### Intracellular target-binding of KMO inhibitors is detected in the TAPS assay

LC-MS/MS detection of the KMO inhibitors (Cpds 1-3) in cell population lysates in the TAPS assay correlated strongly and in a concentration dependent way with levels of target expression, indicating specific inhibitor affinity for KMO (Figure 3a). MS detection of inactive compound 4 demonstrated very low and approximately equal signal in all cell population lysates, indicating little or no specific affinity for KMO (Figure 3a). Furthermore, LC-MS/MS compound detection in fluorescent KMO-positive samples reflected the half maximal inhibitory concentrations generated in the KMO activity assay (Figure 3b). This suggests that the compounds are demonstrating intracellular affinity for KMO and not its E2-Crimson fusion partner. These results show that the TAPS assay protocol can be utilised to identify cell permeable compounds with specific binding affinity for KMO.

### Intracellular target-binding compounds can be detected from a pooled compound mixture

To test the multiplexing potential of the technique, 103 compounds at a final concentration of 20 μM each, including validation compounds 1, 2 and 3, were assayed simultaneously in one cellular incubation mix. Compounds 1, 2 and 3 were the only compounds to be detected solely in the high fluorescence-KMO positive sample indicating that the TAPS assay can be applied for testing medium sized compound collections (Figure 3b). This screen also identified compounds with equal detection across all samples indicating non-specific binding or high cellular permeability. These results were verified by compound testing in the KMO activity assay (Figure 3b). Binding specificity was shown to correlate with inhibitory activity in the activity assay confirming reliable identification of hit and false positive compounds in the TAPS assay.

**Figure 3.**
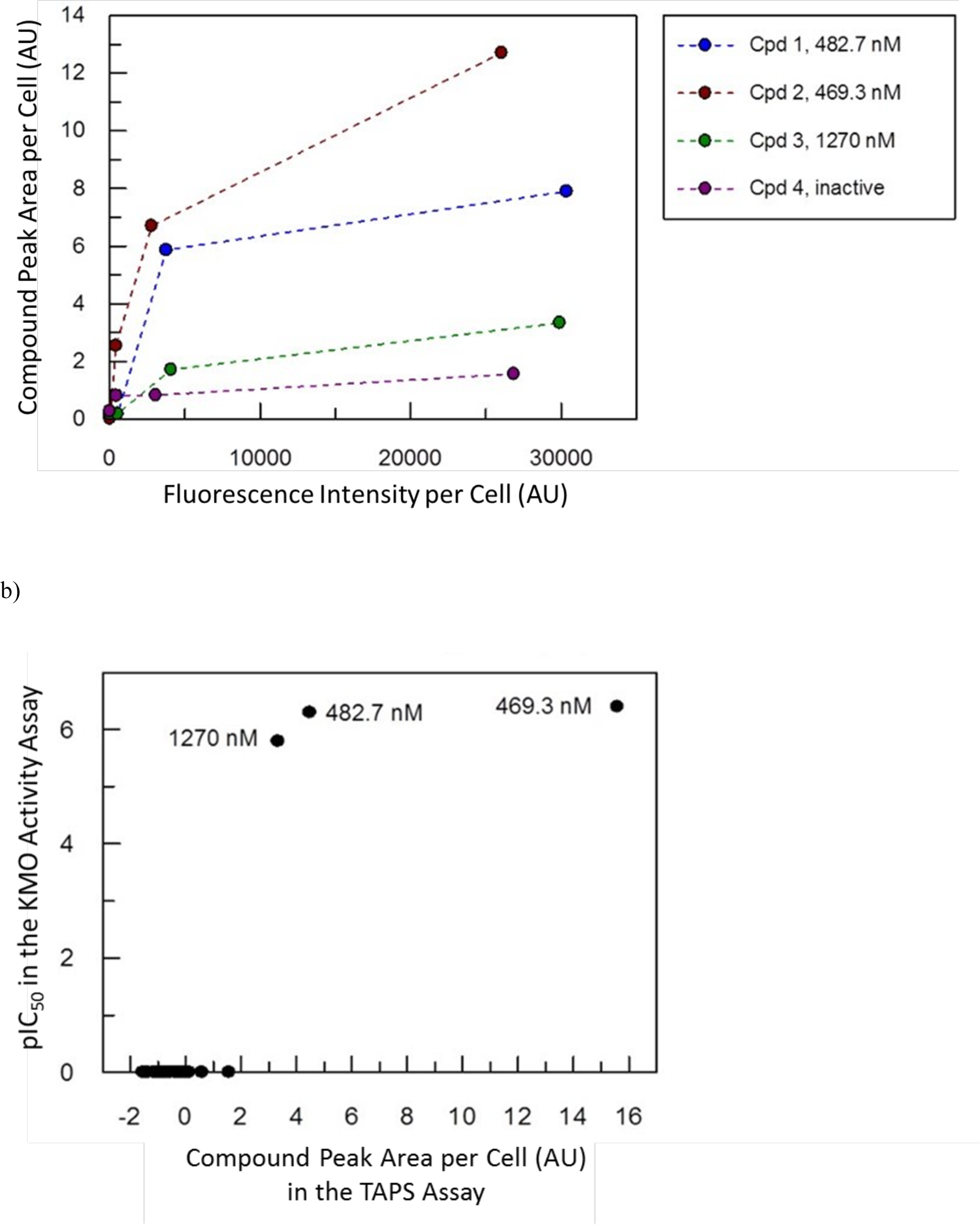
a) The quantity of compound detected by MS for known KMO inhibitors (compounds 1 – 3), represented here as the total peak area detected in a sorted cell sample divided by the number of cells in the population to give compound peak area per cell, increases with the expression of fluorescent target protein detected by FACS. The quantities of validation compounds 1 – 4 detected by MS in the TAPS assay also correlate with the inhibitory activities of each compound, b) Plot showing the correlation between inhibitory activity determined in the KMO activity assay and the quantity of compound detected in cells in the “high” fluorescence/KMO population in the TAPS assay for the compounds in the pooled screen. Each • represents one compound in the screen. Cpds 1 -3 are indicated by their IC_50_ values.

## Discussion

Monitoring of drug-target engagement inside cells represents a challenge in the early drug discovery process. Molina *et al* ^[9]^ described the cellular thermal shift assay (CETSA) which addresses intracellular drug-target engagement but also confirmed that the method is unsuitable for inhomogenous proteins or those which do not aggregate upon unfolding of the ligand-binding domain. Thus there is a need for methods which are applicable to difficult target proteins and for simultaneously testing large sets of test compounds for the evaluation of drug-to-protein binding inside cells. The TAPS assay compares compound detection in cells which demonstrate variable expression of target protein. Utilising known inhibitors of our selected target enzyme, it was shown that the specific affinity of these compounds for the target could be indicated by comparing compound accumulation in cells with over-expressed levels of target protein compared to non-specific accumulation in normal cells. Compounds with equivalent accumulation in overexpressing cells compared to untransfected cells are identified as non-specific binders and excluded. Because only cell permeable target specific binders are comparatively enriched in overexpressing cells, relative compound binding affinity and cellular permeability can be revealed simultaneously by MS analysis. Key control experiments include testing the compound(s) of interest in incubation with variably and transiently transfected cells, fluorescent protein only transfected cells, and, if needed, relevant target mutants and related proteins.

We selected KMO, a challenging but increasingly important target enzyme, for development of this assay. Protein targets, like KMO, which require complexation with cellular interactome proteins or chemical co-factors pose difficulties for *in vitro* biochemical preparation and testing. To overcome these specific challenges we omitted enzyme purification steps and assayed target enzymes directly in mammalian cells in the presence of co-factors and intracellular binding proteins required for enzyme functionality. Successful application of this method to membrane proteins indicates suitability for assay of other challenging targets such as full length protein-protein interaction partners or proteins which are only stable in the presence of nucleic acids or lipids.

KMO was detectable in the cellular matrix following fusion with E2-Crimson^[20]^. FACS and confocal imaging experiments confirmed that cellular E2-Crimson fluorescence reliably indicated KMO expression and functionality. Transient transfection of cells inevitably resulted in variable expression of fluorescent target protein allowing extraction of a negative control sample and a KMO-positive sample from the same cellular drug incubation. This step in the assay facilitates discrimination between compounds demonstrating non-specific cellular component binding in all cells from compounds exhibiting specific binding affinity in cells expressing higher concentrations of target protein. Detection of the known KMO inhibitors in the TAPS assay correlated in a concentration dependent manner with levels of target expression, indicating specific affinity for KMO. The known inactive compound demonstrated an invariable signal regardless of target expression, indicating little or no specific affinity for KMO but also no unspecific or promiscuous binding cellular proteins. Detection of these compounds in the TAPS assay reflected the half maximal inhibitory concentrations generated using the KMO activity assay. Differences in the apparent intracellular binding curves, particularly of the two compounds with similar IC_50_s are most likely connected to cellular uptake, unspecific binding events or compound-specific ionisation during MS readout.

As a next step in the TAPS assay development we tested the multiplexing potential of the technique. 103 compounds were assayed simultaneously in one cellular incubation mix. Detection of the three known KMO inhibitors in the high fluorescence-KMO positive sample suggested suitability of TAPS for testing medium sized compound collections. The screen discriminated compounds showing non-specific binding or high cellular permeability from known KMO-binding compounds with results successfully validated in the KMO activity assay. By comparing the output from each method we were able to confirm reliable identification of hit and false positive compounds in the TAPS assay.

The TAPS method is a multi-parameter tool that delivers on several aspects of compound characterisation. Whilst target binding and cellular permeability is assessed for compounds detected by MS readout, compounds demonstrating cellular cytotoxicity are excluded at the FACS step of the assay. FACS analysis was programmed to incorporate only live healthy cells, by utilising forward scatter (FSC) and side scatter (SSC) gating to eliminate dead cells, meaning that compounds with undesirable cytotoxicity potential are eradicated from the screen. Should further activity data be required for compounds tested by TAPS, lysates resulting from FACS analysis can be directly applied to secondary activity assays, such as the enzyme activity assay demonstrated here. In an alternative assay configuration, a cellular permeabilisation step, like automated optoinjection using LEAP, can successfully be incorporated in the TAPS assay to enable utilisation of the TAPS method for compounds which are not required to permeate the cells (data not shown).

We currently consider that the TAPS assay could be quickly and smoothly integrated into biochemical and biophysical affinity selection, enzymatic and phenotypic screening workflows to prove that any hit compounds engage with the target under investigation. In addition, the TAPS assay might also offer significant value in its potential for medium throughput, multiplexed screening of experimentally challenging protein targets, like intracellular membrane proteins, which are otherwise hindered by a lack of a suitable screening technique. Development and application of the TAPS assay for screening compounds against KMO inside cells indicates that this technique is suitable for targeting other challenging proteins. Besides screening intracellular membrane proteins we also consider TAPS to be the method of choice for intracellular affinity selection assays for larger scale multi-protein complexes containing targetable protein-protein interactions or not easily purifiable targets, such as full length kinases. The excellent quantification possibility of both, protein concentration via fluorescent protein intensity, as well as compound concentration via MS will also allow addressing so far unresolved questions about inhibitor to protein stoichiometry in disease versus healthy cells. An other further iteration of this method will utilise primary tissue cultures. This will enable optimal, physiologically relevant host cell selection tailored to specific target proteins. This may prove especially relevant in cancer therapy since the assay could be applied to drug screening in biopsied tumour cells.

## Conclusion

In conclusion, we report a novel cellular screening technique for assessing intracellular drugtarget engagement. TAPS is a dual parameter technique which is multiplexed with respect to compound and, in a single assay method, enables identification of compounds which are cell permeable, have low cytotoxicity and demonstrate specific affinity for the target protein in a physiologically relevant environment. We demonstrated the potential for the TAPS assay by successful screening of the poorly soluble and challenging protein human, KMO. With target validation and target engagement representing two key limiting factors within the revived importance of phenotypic and high-content screening, we present this solution, which uses off-the-shelf equipment accessible to many scientists in the field. With equally high value, this straightforward workflow will allow direct screening of new targets by affinity selection that until now have only been accessible by indirect detection. We regard important potential targets to include intercompartmental membrane proteins including nuclear envelope transmembrane proteins known to be involved in many cancers. Drug discovery processes are often hindered by issues encountered during preparation of difficult target proteins. Screening of target proteins in a physiologically relevant environment using this method may present a solution, as demonstrated by successful screening of the poorly soluble and challenging protein human KMO.

## References

1. Durham, T.B., Blanco, M.J. Target engagement in lead generation, Bioorg. Med Chem. Lett. 15(5) (2015): 998–1008

2. Kenakin, T., Bylund, D.B., Toews, M.L., Mullane, K., Winquist, R.J., Williams, M. Replicated, reliable and relevant-target engagement and pharmacological experimentation in the 21^st^ century. Biochem Pharmacol. 87(1) (2014): 64–77.

3. Schurmann, M., Janning, P., Ziegler, S., Waldmann, H. Small molecule target engagement in cells. Cell Chem Biol. 23(4) (2016):435–41.

4. Vinegoni, C., Dubach, J.M., Feruglio, P.F., Weissleder, R. Two-photon Fluorescence Anisotropy Microscopy for Imaging and Direct Measurement of Intracellular Drug Target Engagement. IEEE J Sel Top Quantum Electron 22(3), 2016.

5. Dubach, J.M., Kim, E., Yang, K., Cuccarese, M., Giedt, R.J., Meimetis, L.G., Vinegoni, C., Weissleder, R. Quantitating drug-target engagement in single cells in vitro and in vivo. Nature Chemical Biology 13 (Feb 2017): DOI:10.1038

6. D’Allessandro, P.L., Buschmann, N., Kaufmann, M., Furet, P., Baysang, F., Brunner, R., Marzinik, A., Vorherr, T., Stachyra, T.M., Ottl, J., Lizos, D.E., Cobos-Correa, A. Biorthogonal Probes for the Study of MDM2-p53 Inhibitors in Cells and Development of High-Content Screening Assays for Drug Discovery. Angew. Chem. Int. Ed. 2016, 55: 16026–16030.

7. Rutkowska, A., Thomson, D.W., Vappiana, J., Werner, T., Mueller, K.M., Dittus, L., Krause, J., Muelbaier, M., Bergamini, G., Bantscheff, M. A Moldular Probe Strategy for Drug Localization, Target Identification and Target Occupancy Measurement on Single Cell Level. ACS Chem. Biol. 2016, 11: 2541–2550.

8. Simon, G.M., Niphakis, M.J., Cravatt, B.F. Determining target engagement in living systems. Nature Chemical Biology 2013, 9: 200–205.

9. Molina, D.M., Jafari, R., Ignatushchenko, M., Seki, T., Larsson, E.A., Dan, C., Sreekumar, L., Cao, Y., Nordlund, P. Monitoring Drug Target Engagement in Cells and Tissues Using the Cellular Thermal Shift Assay. Science 341, 84 (2013): 84–91.

10. Jafari, R., Almqvist, H., Axelsson, H., Ignatushchenko, M., Lundback, T., Nordlund, P., Martinez Molina, D. The cellular thermal shift assay for evaluating drug target interactions in cells. Nat Protoc. 9(9) (2014): 2100–22.

11. Martinez Molina, D., Nordlund, P. The cellular thermal shift assay: a novel biophysical assay for in situ drug target engagement and mechanistic biomarker studies. Annu Rev Pharmacol Toxicol. 56 (2015): 141–61.

12. Giorgini, F., Guidetti, P., Nguyen, Q.V., Bennett, S.C., Muchowski, P.J. A genomic screen in yeast implicates kynurenine 3-monooxygenase as a therapeutic target for Huntington’s disease. Nat Genet. 2005 May; 37(5):526–531.

13. Mole DJ, Webster S, Uings I, Zheng X, Binnie M, Wilson K et al. Kynurenine–3–monooxygenase inhibition prevents multiple organ failure in rodent models of acute pancreatitis. Nat Med 2016; 22: 202–209.

14. Okamoto, H., Yamamoto, S., Nozaki, M., and Hayaishi, O. (1967) On the submitochondrial localization of L-kynurenine-3-hydroxylase. Biochem. Biophys. Res. Commun. 26, 309–314

15. Uemura, T. & Hirai, K. L-Kynurenine 3-Monooxygenase from Mitochondrial Outer Membrane of Pig Liver: Purification, Some Properties, and Monoclonal Antibodies Directed to the Enzyme. J. Biochem. 123, 253–262 (1998).

16. Wilson, K., Mole, D.J., Binnie, M., Homer, N.Z., Zheng, X., Yard, B.A., Iredale, J.P., Auer, M., Webster, S.P. Bacterial Expression of human kynurenine 3-monooxygenase: solubility, activity, purification. Protein Expr Purif. 95 (2014): 96–103.

17. Wilson K, Mole DJ, Homer NZM, Iredale JP, Auer M, & Webster SP. A Magnetic Bead-Based Ligand Binding Assay to Facilitate Human Kynurenine 3-Monooxygenase Drug Discovery. Journal of Biomolecular Screening (2015) 20(2): 292–8.

18. Pellicciali, R., B. Natalini, et al. (1994). “Modulation of the kynurenine pathway in search for new neuroprotective agents. Synthesis and preliminary evaluation of (m-nitrobenzoyl)alanine, a potent inhibitor of kynurenine-3-hydroxylase.” <u>Journal of medicinal chemistry</u> 37(5): 647–655.

19. Rover, S., A. M. Cesura, et al. (1997). “Synthesis and biochemical evaluation of N-(4-phenylthiazol-2-yl)benzenesulfonamides as high-affinity inhibitors of kynurenine 3-hydroxylase.” Journal of medicinal chemistry 40(26): 4378–4385.

20. Strack, R.L., Hein, B., Bhattacharyya, D., Hell, S.W., Keenan, R.J., Glick, B.S. A Rapidly Maturing Far-Red Derivative of DsRed-Express2 for Whole-Cell Labeling. Biochemistry (2009) 48(35): 8279–8281.

